## Supplementary material for "Deep learning classification of lipid droplets in quantitative phase images": Supporting Information - FINAL.docx

**Supplementary Figure and File**

**Supplementary Figure 1** – The supervised machine learning steps used to train a non-deep, non-convolutional classifier to label pixels corresponding to subcellular lipid droplets.

**Supplementary File 2**: Feature importance is easily determined by evaluating the average relative position of each feature across all decision trees in either the random forest or XGBoost methods. Features closer to the root of the trees are more important to overall classification decision. This spreadsheet describes the relative importance of the 80 extracted features from a trained random forest model, describes the image filtering functions use for feature extraction including their parameterization, and defines which Python libraries were used. Model transparency and interpretability is a key advantage of decision-tree based methods such as random forest or XGBoost. Other methods, such as neural networks, are often impossible to interpret or understand.
