## Supplementary figures and images for "Deep learning classification of lipid droplets in quantitative phase images"

### Supporting Information FIGURE 1.png

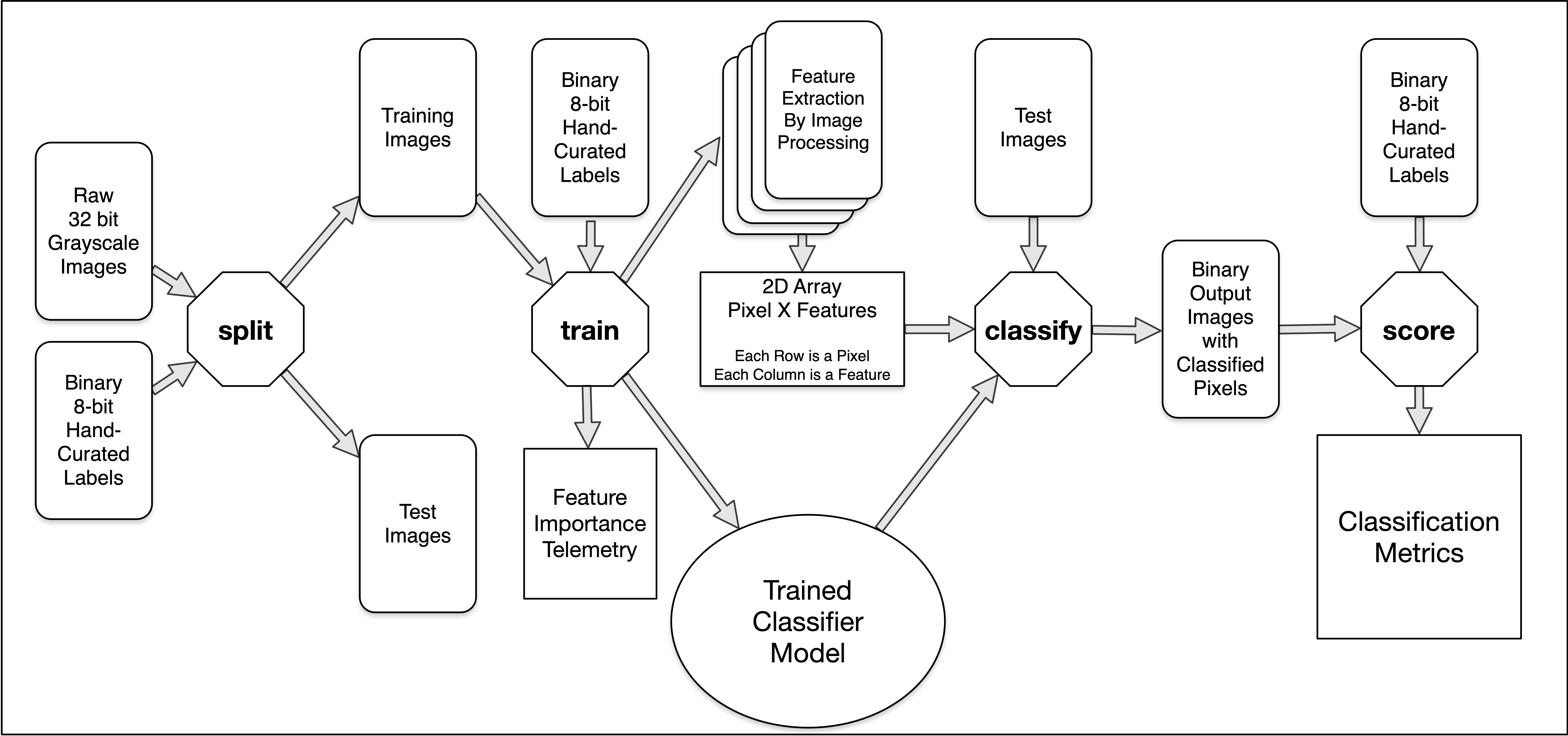
